# Sensitive determination of proteolytic proteoforms in limited microscale proteome samples

**DOI:** 10.1101/566109

**Authors:** Samuel S.H. Weng, Fatih Demir, Enes K. Ergin, Sabrina Dirnberger, Anuli Uzozie, Domenic Tuscher, Lorenz Nierves, Janice Tsui, Pitter F. Huesgen, Philipp F. Lange

## Abstract

Protein N-termini reveal fundamental regulatory mechanisms and their perturbation in disease. Current terminome identification approaches are limited to whole organs or expandable cultured cells. We present a robust, sensitive, scalable and automatable method for system-wide identification of thousands of N-termini from minute samples. Identification of distinct N-terminal profiles in sorted immune cells, subcellular compartments, clinical biopsies, plasma from pediatric cancer patients, and protease substrates in *Arabidopsis* seedlings demonstrate broad applicability.

## Main text

Protein N termini define different proteoforms arising from limited proteolytic processing, alternative translation initiation and co- or post-translational N-terminal modification^1^. De-regulated proteolytic processing of proteins is a well-known driver of disease resulting in aberrant activation, inactivation or change in function, stability or localization of the protein^2^. Consequently, proteases are considered promising drug targets^3^ and proteolytic proteoforms are used as clinical biomarkers^4^. However, proteolytic and otherwise truncated proteoforms are mostly invisible to current bottom-up proteomics workflows^5^. Methods for the selective enrichment of N-terminal peptides have overcome this limitation and enabled, for example, proteome-wide identification of proteolytic proteoforms *in vitro* from apoptotic cells^6^ and *in vivo* using animal models of kidney disease^7^. Further applications include unbiased discovery of protease substrates^8^, identification of alternative protein translation sites^9^ and characterization of protein N-terminal modifications such as acetylation or myristoylation^10^. Yet, requirements for several hundred micrograms to milligrams of purified proteome as starting material^11^ has restricted N terminome analysis to whole organs or expandable cultured cells. The most sensitive protocol to date enables N termini enrichment from 40µg of 10 pooled purified and isobarically labeled proteomes obtained from milligrams of cultured cell lysate^12^. In contrast, proteome^13^ and phosphoproteome^14^ sample processing have been improved and automated, enabling comprehensive analyses of cell-type specific processes and clinically relevant microscale samples (<20µg). To enable a similar leap in utility for the analysis of proteolytic proteoforms, we developed High-efficiency Undecanal-based N Termini EnRichment (HUNTER), an automatable workflow for the sensitive enrichment of N-terminal peptides from as little as 2µg protein lysate of any cell or tissue using off-the-shelf reagents (Fig. 1a).

**Fig. 1.**
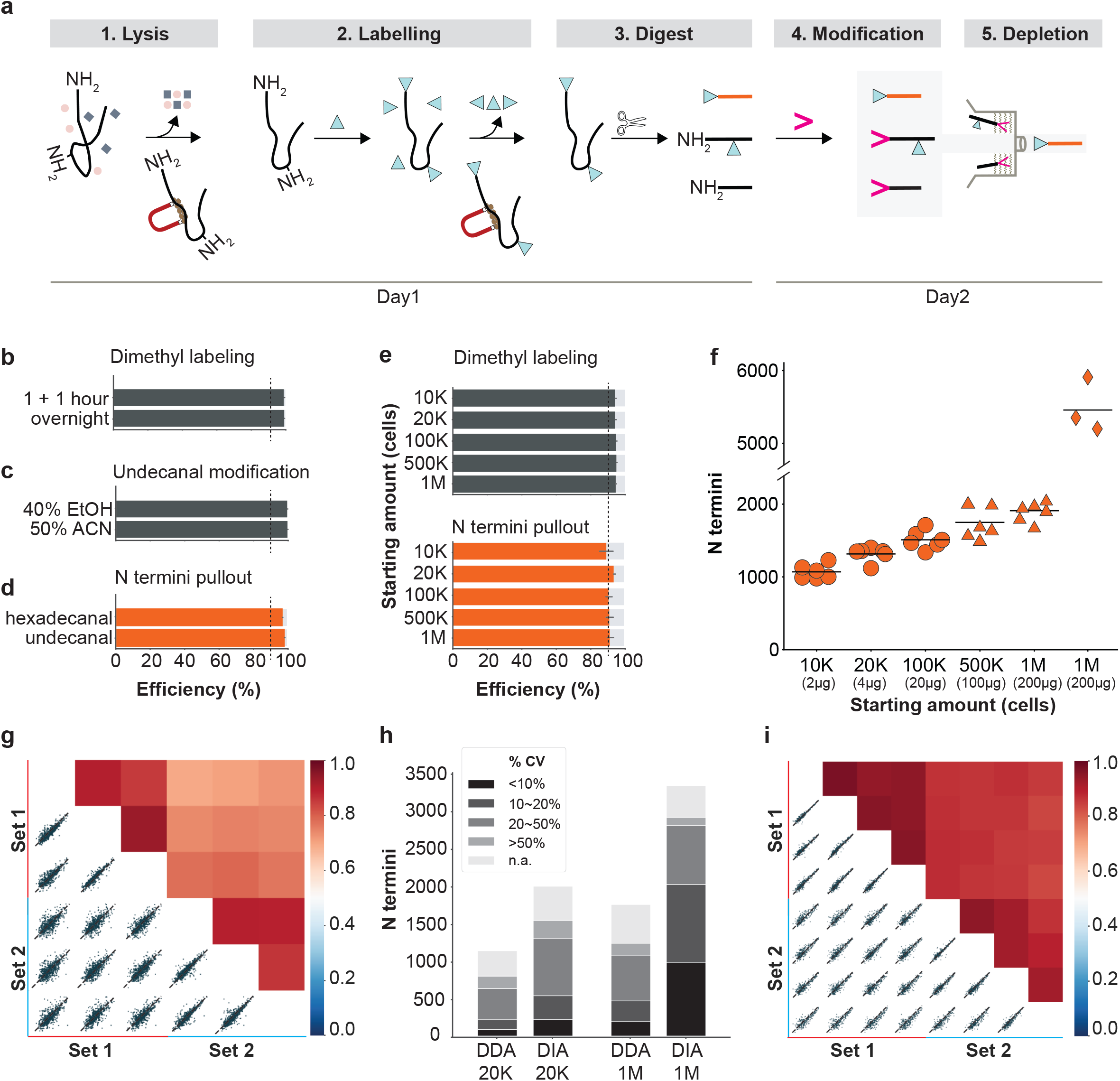
A protocol for sensitive enrichment and identification of N-terminal peptides. (**a**) Workflow: (1) Lysis and protein purification with SP3 magnetic beads; (2) modification of N-terminal α- and lysine ε-amines; (3) proteome digestion; (4) modification of digestion-generated peptide α-amines with undecanal in 40% ethanol (grey background); (5) retention of undecanal-modified peptides on reverse-phase columns, while N-terminal peptides pass through. (**b**-**d**) Optimization of individual steps. (**b**) Completeness of on-bead dimethyl labeling of HeLa cell lysates (n=3 technical replicates). (**c**) Degree of HeLa-derived peptide undecanal modification in organic solvents, assessed by reactivity of unlabeled amines (n=3 technical replicates). (**d**) Comparison of modification reagents-hexadecanal and undecanal-assessed by N termini enrichment from *Arabidopsis thaliana* leaf proteome (n=3 biological replicates). (**e**-**g**) Evaluation of HUNTER workflow using varying starting amounts of HeLa cells. (**e**) Dimethyl labeling efficiency assessed by lysine side chain modification before depletion (top) and N termini enrichment efficiency after enrichment (bottom). (**f**) Number of N termini identified from 10,000 to 1,000,000 HeLa cells. Circles, samples with <1µg peptide analyzed in a single injection; triangles, larger samples with 1µg enriched peptides analyzed per replica; diamonds, with offline high-pH pre-fractionation. Datapoints represent technical replicates, lines indicate the mean. (**g**) Intensity-based Pearson-correlation between two manually processed N termini enrichment datasets from 20,000 HeLa cells, identified by data-dependence acquisition (DDA, each set n=3 technical replicates). (**h**) Comparison of coefficients of variation (CV) for N termini from 20K and 1M HeLa cell starting material quantified by label free DDA or DIA. Average CV of pairwise comparisons of quantified terminal peptides in triplicate analysis is displayed. NAN denotes termini quantified in only one replica. (**i**) Intensity-based Pearson-correlation between two automatically processed sets (each n=4 technical replicates) from human plasma samples, acquired by DDA. All error bars indicate SD.

The most sensitive protocols established to date enrich protein N termini by negative selection^11^, where protein amines are blocked with amine-reactive reagents before proteome digestion. This in turn generates new peptide-N-terminal α-amines that are then exploited for depletion. Unspecific losses and low reproducibility mainly occur in three critical steps (Fig. 1a), during removal of free amino acids and other interfering compounds from the protein lysate, removal of amine-reactive labeling reagents prior to digestion and selective depletion of proteome digestion-generated non-N-terminal peptides (Fig. 1a). The first two purification steps are commonly achieved by protein precipitation and resuspension, resulting in high protein loss and poor reproducibility, particularly in microscale samples^15^. To overcome this, we replaced precipitation-based protein purification by reversible high-efficiency binding to hydroxylated magnetic beads as used in the Single-Pot Solid-Phase-enhanced Sample Preparation (SP3) method^15^. We first established compatibility of SP3 with protein-level dimethyl labeling and found that within 2 hours >99% of primary amines on proteins were successfully blocked from subsequent reaction (Fig. 1b). The third loss-intensive step is depletion, where considerable losses occur due to unspecific binding of dilute N-terminal peptides to surfaces of filters, beads and other consumables (Fig. 1a). Here we adapted a strategy based on attaching hydrophobic hexadecanal to the free peptide a-amines generated by the proteome digest^16^. This increased the hydrophobicity of tryptic non-N-terminal peptides, allowing their retention on a reverse phase liquid chromatography column, while N-terminal peptides were eluted and directly analyzed by MS/MS at acetonitrile concentrations <80%. However, hexadecanal-containing reactions solidified at room temperature and underwent phase separation resulting in losses and lowered reproducibility. We tested the shorter-chain undecanal, which is liquid at room temperature. After optimizing reaction time (Supplementary Fig. S1a,b) and solvent conditions (Fig. 1c), we found that reaction in 40% ethanol for 60 min at 37°C, followed by passing the reaction mixture through commercial C18 reverse phase resins (Supplementary Fig. 1c) allowed direct enrichment with minimal loss of N-terminal peptides (Supplementary Fig. 1d). Additionally, depletion of shorter-chain undecanal-tagged peptides was equally or more efficient compared to hexadecanal (Fig. 1d, Supplementary Fig. 2a,b), resulting in enrichment of N-terminally modified peptides from baseline levels of <10% to >92% after enrichment (Supplementary Fig. 2c). The enrichment efficiency was independent from the amount and source of digested proteome used for pullout, including *Arabidopsis thaliana* leaf and rat brain proteomes (Supplementary Fig. 2d,e).

After optimizing SP3-labeling and undecanal-mediated depletion individually, we evaluated the performance of the combined workflow in a one-pot reaction from lysis to cleanup. Across a wide range of starting material from 1 million HeLa cells, equivalent to 200µg protein lysate, down to as few as 10,000 HeLa cells, or 2µg protein lysate, we observed >94% dimethyl-modified lysine residues and enrichment efficiencies >90% (Fig. 1e). Analysis with an Orbitrap Q-Exactive HF mass spectrometer identified an average of 992, 1,230 and 1,454 N-terminal peptides from 2µg, 4µg and 20µg, respectively, within one hour. For larger samples, only 1µg of the recovered N-terminal peptides were injected, resulting in the identification of 1810 N-terminal peptides on average. High-pH fractionation of N-terminal peptides enriched from 200µg HeLa proteome readily increased this to identification of >5,000 N-terminal peptides (Fig. 1f). Data independent acquisition (DIA) using a spectral library extracted from these fractions further boosted the number of N-termini identified from 200µg starting material in a single 1-hour analysis by 50% to 2,877 (Supplementary Fig. S3a). The reproducibility was similar across the range of starting material, with Pearson correlation factors of 0.89 between manually pipetted replica of 4µg HeLa material and 0.74 between days (Fig. 1g, Supplementary Fig. S3b-e), as reported for label-free single-peptide quantification with minimal starting amounts^14^. DIA analysis of 4µg and 200µg HeLa lysate showed similar correlation coefficients of 0.91 between manually pipetted replica (Supplementary Fig. S3f,g), but markedly improved quantitative precision. Using DIA analysis, 988 N termini were quantified in 200ug HeLa lysate with single peptide CVs <10%, whereas DDA analysis only quantified 192 N termini with CVs < 10% (Fig. 1h). Finally, we established HUNTER on a basic liquid handling system in 96 well format to enable high-throughput sample analysis and to reduce variation from manual pipetting. Automated enrichment of protein N termini from 1 to 5µl of human plasma in four technical replicates achieved an improved intra-assay Pearson correlation of 0.93. The inter-assay Pearson correlation for automated assays on different days and different chromatography columns was 0.86 on average (Fig. 1i).

We next set out to explore the performance and utility of the HUNTER protocol with samples that had so far not been amenable to N terminome characterization. We first analyzed 3 µl non-depleted blood plasma (BP) and aspirated bone marrow interstitial fluid (BM) from three pediatric B-cell acute lymphoblastic leukemia (B-ALL) patients before and after induction chemotherapy. This analysis identified 600 N termini of 244 proteins across all patients, with more low-abundance plasma proteins^17^ identified after N termini enrichment than in a standard proteome analysis (Fig. 2a). Quantitation revealed pronounced differences between plasma and BM, and before and after treatment (Supplementary Fig. S4). Notably, N termini matching complement protein activation sites^18^ decreased in both plasma and bone marrow during chemotherapy (Fig. 2b, Supplementary Fig. 5), in line with the chemotherapy induced complement defects observed in ALL^19^.

**Fig. 2.**
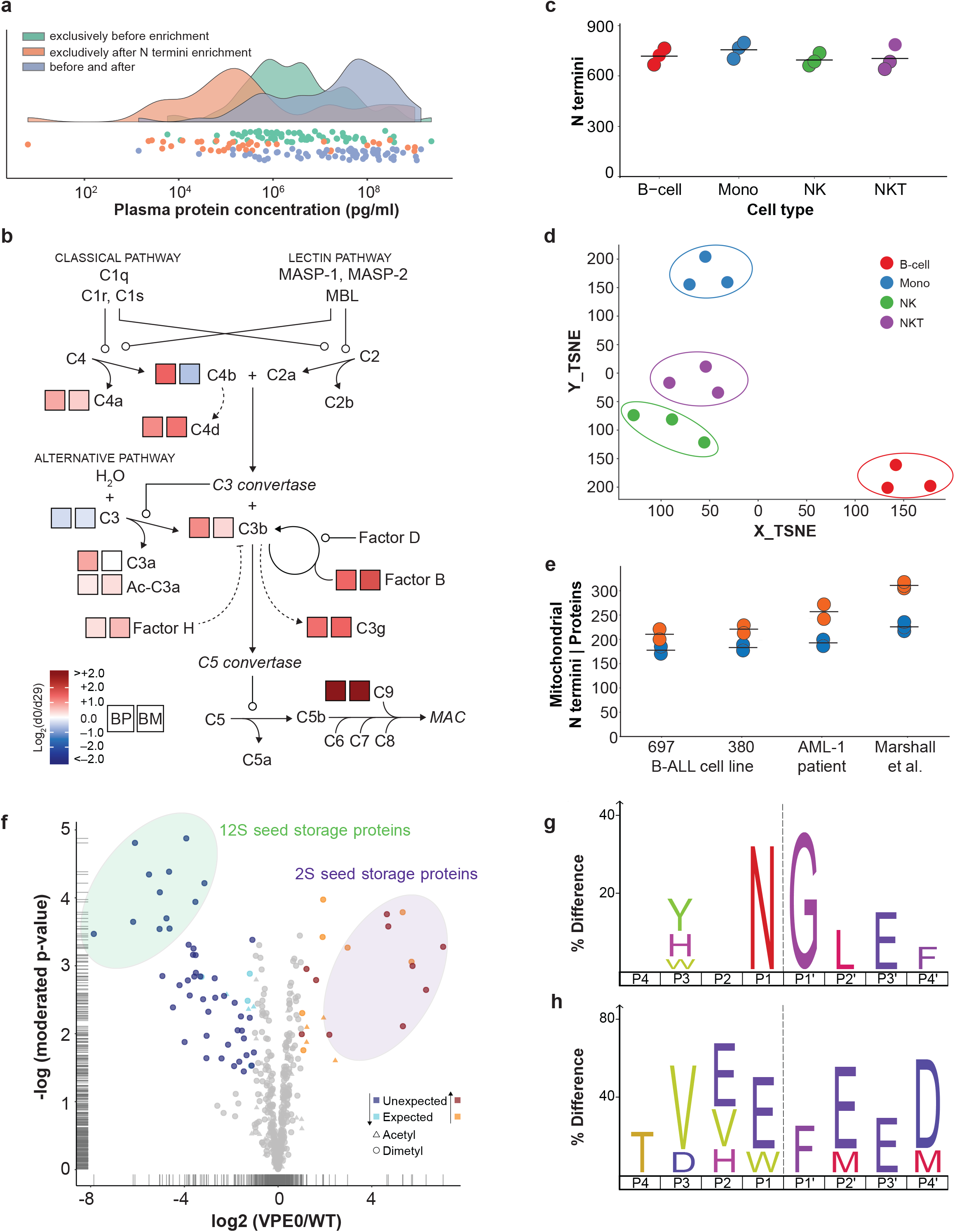
N terminome analysis in microscale samples. (**a,b**) Quantitative analysis of plasma (BP) and bone marrow interstitial fluid (BM) from three B-ALL patients. (**a**) Proteins identified exclusively in plasma before or after N termini enrichment, or in both preparations, mapped to known concentrations. (**b**) Quantification of complement pathway protein N termini. (**c**,**d**) N terminome analysis of sorted peripheral blood mononuclear cells from three healthy donors. (**c**) N termini identified from 30,000 B-cells, monocytes, natural killer (NK) cells and NKT-cells, mean indicated by line. (**d**) Unsupervised t-distributed stochastic neighbor embedding (t-SNE) based on raw log_10_ transformed N terminal peptide intensities. (**e**) Mitochondrial protein N termini enriched from 2.5 million B-ALL 697, B-ALL 380 and AML patient cells compared to reported N termini identification from 12-15 million HeLa cells. Orange, N termini; Blue, proteins. (**f**-**h**) Comparative N terminome analysis of 2.5-day old *Arabidopsis* VPE0 quadruple mutants and WT seedlings. (**f**) Volcano plot. Circle, dimethylated N termini; triangle, acetylated N termini; light blue/orange, significantly (p-value <0.05, log_2_(VPE0/WT)<-1 or>1) less/more abundant N termini matching known or predicted N-termini; dark blue/dark red, significantly (p-value <0.05, log_2_(VPE0/WT)<-1 or>1) less/more abundant N termini matching “unexpected” positions within the protein models. (**g**) iceLogo representing 24 cleavage sites deduced from 47 unexpected N-terminal peptides significantly depleted in the VPE0 mutant filtered for redundancy created by exopeptidase ragging. (**h**) iceLogo representing 7 filtered cleavage sites deduced from 10 unexpected N-terminal peptides significantly more abundant in the VPE0 mutant. Dashed line indicates cleavage site.

We then explored differences in protein maturation between four different peripheral blood monocyte populations from healthy donors isolated by fluorescence-activated cell sorting. For each population we analyzed triplicates of 30,000 sorted cells and identified between 646 and 803 N termini (Fig. 2c). Unsupervised dimensionality reduction based on N termini abundance clearly separated the different cell types, with replica of each cell type grouped in close proximity (Fig. 2d). Next, we asked if HUNTER could support investigation of proteolytic processes in subcellular compartments of limited samples, such as pediatric patient biopsies. This type of study was previously restricted to cultured cells of which large quantities of source material could be obtained. We first optimized a crude subcellular fractionation of mitochondria using mild pressure cycling assisted cell lysis of 2.5 million cells in 30µl buffer, resulting in a threefold increase of N termini originating from known mitochondrial proteins (Supplementary Fig. 6a). HUNTER applied to mitochondrial fractions from less than 2.5 million Acute Myeloid Leukemia (AML) blasts obtained by bone marrow aspiration from a pediatric patient enabled detection of 233 N termini for 181 mitochondrial proteins, with similar numbers identified in two B-ALL cell lines (Supplementary Fig. 6b,c). Compared to a recent study of mitochondrial protein processing in human cells^20^, HUNTER identified on average 73% of the protein termini and 81% of proteins from only about 1/10th of the starting material (Fig. 2e).

Finally, we tested the utility of HUNTER for protease substrate profiling in small specimen by comparing *A. thaliana* wild type seedlings with a quadruple knock-out of all four genes coding for vacuolar processing enzymes (VPEs) resulting in altered seed storage protein processing^21^. HUNTER analysis of single seedlings in combination with duplex stable-isotope labeling showed high biological variation in early development (Supplementary Fig. S7). To reduce biological variability and increase coverage, we pooled three seedlings per condition (Supplementary Fig. S8) and found that 75 of 933 quantified N termini showed significant changes between both lines (Fig. 2f). 54 N termini more abundant in wild-type mostly reflected altered processing of 12S seed storage proteins and predominantly matched the known VPE sequence specificity for cleavage after Asn (Fig. 2g). The 21 N-terminal peptides with increased abundance in the VPE null mutant indicated alternative processing of 2S seed storage proteins, preferentially between Glu or Trp and Phe (Fig. 2h), and increased activation of cathepsin B3 and the germination-specific cysteine proteases CP1 that might help to compensate for the lack of VPEs (Supplementary Fig. S8).

In summary, HUNTER is a highly sensitive, universal and scalable protocol for enrichment of protein N termini from crude protein lysates. HUNTER is well suited for automation even on basic liquid handling systems, as it is based on standard magnetic bead and cartridge technology, does not require protein precipitation and avoids phase separations. We have shown successful application in systems as diverse as rat brain and plant leaf tissue, human plasma, sorted peripheral blood cell populations, subcellular fractions enriched for mitochondria and individual *A. thaliana* seedlings. With sensitive identification and reproducible quantification of >1,000 protein termini from starting amounts of as little as 10,000 HeLa cells or 2 µg of protein lysate and >5,000 termini from 200µg of protein lysate, HUNTER enables comprehensive analysis of proteolytic processes and protein N-terminal modifications in microscale samples from a wide range of precious limited biological samples and clinical biopsies.

## Methods

### Cell culture and human samples

HeLa cells (American Type Culture Collection; cat. no. CCL-2) were cultured in RPMI 1640 medium (ThermoFisher Scientific; cat. no. 11875-093) with 10% Cosmic Calf Serum (GE Healthcare Life Sciences; cat. no. SH30087.04) and maintained in a humidified incubator at 37°C with 5% CO2. Cultured cells were collected using 0.25% Trypsin-EDTA (ThermoFisher Scientific; cat. no. 25200056), centrifuged at 800 x g and washed with PBS (ThermoFisher Scientific; cat. no. 10010023) to collect pellets of different cell quantities. Cell pellets were frozen and stored in −80°C freezer until further lysis. B-cell acute lymphoblastic leukemia (B-ALL) cell lines 380 (ACC 39) and 697 (ACC 42) cells were procured from DSMZ (Braunschweig, Germany). B-ALL cell lines were cultured in RPMI 1640 media supplemented with 10% heat-inactivated fetal bovine serum (ThermoFisher Scientific; cat. no. 10082147) and 2 mM L-Glutamine (ThermoFisher Scientific; cat. no. 25030081) and maintained at 37 °C in 5% CO_2_. Commercial human blood plasma was purchased from STEMCELL Technologies (cat. no. 70039). Primary pediatric B-ALL and AML patient mononuclear cells enriched from bone marrow aspirates, plasma (BP) and bone marrow interstitial fluid (BM) were retrospectively sourced from the Biobank at BC Children’s Hospital (BCCH) following informed consent and approval by the University of British Columbia Children’s and Women’s Research Ethics Board (REB #H15-01994) in agreement with the Declaration of Helsinki. Patient BP and BM samples were collected at the time of diagnosis (D0) and 29 days after induction chemotherapy (D29). Peripheral blood mononuclear cells (PBMC) from healthy donors were obtained following informed consent and approval by the University of British Columbia Children’s and Women’s Research Ethics Board (REB #H10-01954). Individual populations were obtained by Fluorescence Activated Cell Sorting using the following antibody combinations: CD19+ for B-cells, CD14+ for monocytes, CD3- CD56+ for natural killer (NK) cells and CD3+ CD56+ for NK T-cells (NKT cells).

### Plant material

*Arabidopsis thaliana* Col-8 wild type (accession N60000) and VPE0 mutant (accession N67918) seed stocks were obtained from the Nottingham Arabidopsis Stock Center (NASC). *A. thaliana* Col-8 plants were stratified for 3d at 4°C and subsequently grown on soil at short day conditions (9h light with an intensity of 100 µE m^-2^ s^-1^ at 22°C and 15 h darkness at 18°C, 75% RH). Leaves of 6-week-old plants were harvested and snap frozen in liquid nitrogen. For seedling experiments, *A.thaliana* seeds were stratified for 3d at 4°C and germinated for 2.5d (5d for single seedling experiment) on filter paper at a short day time regime (9h light with an intensity of 110 µE m^-2^ s^-1^ at 22 °C and 15 h darkness at 18 °C).

### Rat brain samples

Rat brains were obtained from Wistar rats that were sacrificed for liver perfusion experiments at the University Hospital Düsseldorf as approved by local authorities (LANUV NRW #G287/15) and immediately snap frozen in liquid nitrogen.

### Preparation of Stage-tips

Four small circular Empore™ SPE C18 disks (Sigma, cat. no. 66883-U) were punched with a flat-end needle (Hamilton, cat. no. 90517). A straightened paper clip was used to gently push down the C18 disks into a P200 pipette tip (VWR, cat. no. 89079-474).

### High-pH reversed phase fractionation

Fractionation was performed with an Agilent 1100 HPLC system equipped with a diode array detector (254, 260, and 280 nm). HPLC system was installed with a Kinetic EVO C18 column (2.1mm×150mm, 1.7µm core shell, 100Å pore size, Phenomenex). The samples were run at a flow rate of 0.2ml per minute using a gradient of mobile phase A (10mM ammonium bicarbonate, pH 8, Fisher Scientific, cat. no. BP2413-500) and mobile phase B (acetonitrile, Sigma-Aldrich, cat. no. 34998-4L) from 3% to 35% B over 60 min. Fractions were collected every minute across the elution window for a total of 48 fractions, then concatenated to a final set of 12 (e.g. fraction 1+13+25+37 as final fraction 1). All the fractions were dried in a SpeedVac centrifuge and resuspended in 0.1% FA in water (Thermo Scientific, cat. no. SC2352911) prior to mass spectrometry analysis.

### High-efficiency Undecanal based N Termini EnRichment (HUNTER)

#### Preparation of *HeLa* and peripheral blood mononuclear sorted cell lysates

The HeLa cell and sorted cell samples were first lysed in 1.5ml protein Lobind tube (Eppendorf; cat. no. 022431081) with lysis buffer consisting of 1% sodium dodecyl sulphate (Fisher BioReagents, cat. no. BP8200-500) and 2X Thermo Halt protease inhibitor cocktail (Thermo Scientific, cat. no. 1861279) in 50 mM HEPES, pH 8.0 (Sigma, cat. no. H4034-1KG). The lysate was heated at 95°C for 5 min, then chilled on ice for another 5 min. Any liquid condensation or droplets was spun down by centrifugation. Benzonase (EMD Millipore; cat. no. 70664-3) was added at a ratio of 1 unit to 37µg of DNA and incubated at 37°C for 30 min. Then DTT (DTT; Fisher BioReagents; cat. no. BP172-25) was added to 10mM and incubated at 37°C for 30 min, followed by addition of 2- chloroacetamide (CAA; Sigma-Aldrich; cat. no. C0267-100G) to 50mM and further incubation at RT in the dark for 30 min. To quench the alkylation, DTT was added to a final concentration of 50 mM and incubate at RT in the dark for 20 min. Protein Lobind tubes were used during all sample handling steps.

#### Preparation of mitochondrial enrichment samples

Mitochondrial enrichment was performed on 2.5 million cells from two B-cell lines (697, 380), and 2.5 million bone marrow monocytes from a pediatric AML patient (AML-1). All samples were processed in technical replicates (n=2 or n=3). Cells in mitochondrial isolation buffer (1 mM EGTA/HEPES pH 7.4, 200 mM Sucrose, 1 x Halt protease inhibitor) were disrupted by Pressure Cycling Technology (PCT) using a Barocycler EXT2320 and a PCT 30µL MicroTube (Pressure BioSciences, Easton, Massachusetts, United States). The cell samples were homogenised and lysed using 15 cycles of 25kpsi for 20 seconds and followed by 20 second at ambient pressure at 26°C. Cells were subsequently centrifuged at 900 x *g* and the pellet fraction (Mitochondrial fraction 1, M1) was collected. The supernatant was transferred to a new tube and centrifuged at 13,000 x *g* to collect the second pellet fraction (Mitochondrial fraction 2, M2) and cytosolic supernatant (cytosolic fraction, C). Pellet fractions M1 and M2 made up the mitochondrial enriched portion. Proteins were reduced and denatured as described for HeLa samples.

#### Preparation of *Arabidopsis thaliana* seedling lysates

Single 5 day-old *Arabidopsis* seedlings, or three 2.5 day-old pooled germinating seeds, were lysed with a buffer consisting of 4% sodium dodecyl sulphate, supplemented with 2X Thermo Halt protease inhibitor cocktail in 100 mM HEPES, pH 7.5 for 10 min at 95 °C. Mechanical disruption was performed with single use pestles in protein Lobind tubes, followed by heating to 95 °C for 10 min and subsequent chilling on ice for 5 min. Proteomes were reduced with 5 mM DTT for 30 min at 56°C, alkylated with 15 mM iodoacetamide (IAA) for 30 min in the dark at RT, and quenched by addition of additional 15 mM DTT and incubation for 15 min at RT.

#### SP3 bead binding and proteome clean up

After reduction and alkylation, prepared SP3 beads were added to protein mixtures with a 1:10 ratio (w/w) protein / SP3 beads. Pure 100% ethanol was added to a final volume 80% v/v to initiate binding. After 18 min incubation at RT, supernatant was removed with assistance of a magnetic stand and the beads were rinsed two times with 400µL 90% ethanol. Beads were resuspended by pipet mixing, with 30s break between each step to allow beads to settle on the magnetic stand. The remaining ethanol was spun down prior to the removal of supernatant and beads were resuspended in 30µL 200mM HEPES, pH 7.0.

#### Protein dimethyl labeling

2M freshly prepared formaldehyde solution (Sigma-Aldrich, cat. no. 252549) and 1M sodium cyanoborohydride (Sigma-Aldrich, cat. no. 296813) were added to 30mM and 15mM final concentration, respectively. In the *Arabidopsis* seedling experiment, ^12^CH_2_O formaldehyde was used for labeling of WT proteome and heavy ^13^CD_2_O formaldehyde (Sigma, cat. no. 596388) for the VPE0 quadruple mutant proteome. The lysate was incubated at 37°C for 1 h in an oven, before repeated addition of fresh labeling reagents and incubation for another hour. To quench the reaction, 4M Tris buffered to pH 6.8 (Fisher BioReagents, cat. no. BP153-1) was added to a final concentration of 600 mM (500 mM for *Arabidopsis* seedling proteome) and incubated at 37°C for 3 hours (30 min for *Arabidopsis* seedling proteome). For removal of excess reagents, new SP3 beads were added at a 1:5 ratio and protein bound by addition of 100% ethanol to a final concentration of 80% v/v ethanol. Beads were settled on a magnetic stand after 15 min incubation at RT, supernatant removed, and the beads washed twice with 400µL of 90% ethanol. The tube was briefly centrifuged to collect and remove the remaining wash solution before resuspension of the beads in 30µL trypsin (1mg/ml, Promega, cat. no. V5113) in 200mM HEPES buffer, pH 8.0. Beads were fully immersed in the solution and the trypsin to protein ratio was at least 1:100. After incubation at 37°C in an oven for at least 13 hours, 10% of the sample was removed to assess dimethyl labeling efficiency or to quantify protein abundance (pre-HUNTER sample). The reaction was mixed by tapping and 30s sonication after addition of each new reagent. Differentially labeled *Arabidopsis* seedling WT and VPE0 proteomes were pairwise combined after this step.

#### Enrichment of protein N termini by undecanal-assisted negative selection

100% ethanol was added to the proteome digest to 40% v/v before addition of undecanal (EMD Millipore, cat. no. 8410150025) at an undecanal/peptide ratio of 20:1 w/w (50:1 for *Arabidopsis* seedling samples) and addition of 1M sodium cyanoborohydride to a final concentration of 30mM. The pH was confirmed between pH 7-8 before incubation at 37°C for 1 hour. The reaction was sonicated in a water bath at 60kHz for 15 seconds and bound to magnetic rack for 1 min. The supernatant was transferred to a new Lobind tube and acidified with 0.5% trifluoroacetic acid (TFA) (Sigma-Aldrich, cat. no. T6508-100ml) in 40% ethanol to pH 3-4 before loading onto a C18 column for removal of undecanal-tagged peptides. Different columns were chosen to provide sufficient binding capacity for excess undecanal reagent: Self-packed 4-layered C18 stage-tips were chosen for 1 to 5µg protein; microspin column (Nest Group Inc, cat. no. SEM SS18V) for 5 to 20µg protein; macrospin column (Nest Group Inc, cat. no. SMM SS18V) for 20 to 100µg protein; sep-pak columns (Waters, cat. no. WAT054960) for 100-1000 µg protein; HR-X (M) spin columns (Macherey-Nagel, cat. no. 730525) for experiments with *Arabidopsis* and rat brain proteome. The sample volumes were topped up with 0.1% TFA in 40% ethanol to a loading volume was 80µL, 200µL, 400µL and 500µL for stage-tips, microspin, HR-X (M) spin, macrospin column, and sep-pak respectively. Before loading the samples, the stage-tips were conditioned with 100µL methanol and followed by 100µL 0.1% TFA in 40% ethanol whereas microspin column, macrospin column, HR-X (M) spin columns and sep-pak were conditioned with a volume of 200µL, 200µL, 400 µL and 700µL respectively. After the conditioning of C18 columns, the samples were then loaded and the flow-through was collected in 1.5ml protein Lobind tubes. The ethanol in the collected flow-through was removed by vacuum supported evaporation, peptides were resuspended in. 0.1% TFA in HPLC water and desalted using home-made C18 stage-tips or commercial reverse-phase C18 spin columns.

### Automated HUNTER

Human peripheral blood plasma (STEMCELL Technologies, cat. no. 70039) and plasma and bone marrow interstitial fluid samples from three pediatric B-ALL patients (B-ALL-1, −2, −3) were processed on an epMotion M5073 automated liquid handling system (Eppendorf) controlled by an EasyCon tablet (Eppendorf). The HUNTER protocol was programed with epBlue Studio (ver. 40.4.0.38). The M5073 was configured with: dispensing tool TS50 (1.0-50µL) and TS1000(40- 1000µL), epT.I.P.S. Motion racks (1.0-50µL and 40-1000µL), epMotion gripper, Thermoadapter for 96-PCR plate (skirted), Alpaqua Magnum FLX 96 magnet plate, Eppendorf rack for 24x safe lock, Twin.tec PCR plate 96 (semi-skirted; max. well volume is 250µL).

The following adaptations to the HUNTER protocol were made to achieve optimal automation: 250-300µg protein (maximum 5µL plasma) was processed. Dimethylation was performed at room temperature, the final concentration of formaldehyde was 35mM, and the final concentration of sodium cyanoborohydride was 15mM. 2 units of benzonase were added to 5µl of plasma. Wash steps were programmed to aspirate 10µL more than the dispense volume to ensure full removal of all wash buggers. During the digestion and undecanal labeling steps, the plate was covered with thermal adhesive sealing film (Diamed Lab Supplies Inc., cat. no. DLAU658-1) and incubated at 37°C. Samples and/or beads were mixed on the heater/shaker at 1500rpm for 2 min. To prevent bubbles forming in tips and ensure uniform dispensing, the aspiration speed was set to 10mm/s. All pipetting steps were programmed to aspirate from bottom and dispense from top. Undecanal and ethanol were combined first before dispensing into each well.

#### Preparation of single-pot solid-phase-enhanced sample preparation (SP3) beads

1:1 v/v ratio of hydrophilic (conc. 10µg/µL; GE Life Sciences; cat. no. 4515-2105-050250) and hydrophobic Sera-Mag SpeedBeads carboxylate-modified magnetic beads (conc. 10µg/µL; GE Life Sciences, cat. no. 6515-2105-050250) were combined in a 1.5ml flex tube (Eppendorf, cat. no. 022364111), then place them on a magnetic stand (Life Technologies, cat. no. 12321D) for removal of supernatant. The beads were washed twice and reconstituted in HPLC water (Fisher Scientific; cat. no. W6-4) and stored at 4°C.

#### Fluorometric and colorimetric protein and peptide measurements

To evaluate labeling, binding and elution efficiencies during protocol optimization, peptide concentration and primary amine reactivity were quantified using the Pierce quantitative fluorometric peptide assay (Thermo Fisher Scientific; cat. no. 23290) and Pierce quantitative colorimetric peptide assay (Thermo Fisher Scientific; cat. no. 23275) following the assay protocols.

#### Optimizing dimethyl labeling of proteins

10µg of reduced and alkylated HeLa protein were used as starting material. 2M fresh formaldehyde and 1M sodium cyanoborohydride were added to 30mM and 15mM final concentration in 200 mM HEPES, pH 7.0. The reaction was incubated at 37°C as indicated. After the first incubation, fresh labeling reagents were added and incubated at 37°C for 1 h. Both LC-MS/MS analysis and the amine-reactive quantitative fluorometric peptide assay were performed to evaluate the dimethyl labeling efficiency.

#### Optimizing undecanal modification of peptide α-amines

Frozen plant leaves or rat brains were homogenized in 6M GuaHCl, 0.1M HEPES pH 7.4, 1 mM DTT, 5 mM EDTA and 1X Thermo Halt Protease inhibitor mix with a Kinematica Polytron PT-2500 for 2x 30s at 18 000 rpm (Kinematica, Luzern, Switzerland). Homogenate was filtrated through Miracloth (Merck, Darmstadt, Germany) and cell debris pelleted at 500 g for 5 min, 4 °C. Supernatants were chloroform/methanol precipitated, resuspended in 1:2 diluted homogenization buffer, reduced by incubation with 5 mM DTT for 30 min at 56 °C and alkylated by addition of 15 mM IAA for 30 min in the dark at RT. The reaction was quenched by addition of additional 15 mM DTT and incubation for 15 min at RT.

Dimethylation was performed at protein level with 20 mM heavy formaldehyde (^13^CD_2_O) and 20 mM sodium cyanoborohydride at 37 °C O/N. The next day, fresh 20 mM formaldehyde and sodium cyanoborohydride were both added for a further dimethylation of 2h at 37 °C. Labeled proteins were purified by chloroform-methanol precipitation and concentration determined using the BCA assay (BioRad). Samples were digested over night at 37 °C with MS-grade trypsin (Serva,) at a 1:100 protease:protein ratio in digestion buffer (0.1M HEPES pH 7.5, 5% ACN, 5 mM CaCl_2_). Digestion was prolonged by addition of fresh MS-grade trypsin at a 1:200 ratio for 2h at 37 °C.

Trypsin-generated peptide a-amines were hydrophobically modified by adding undecanal in a 50:1 (w/w) ratio undecanal/proteome and 20 mM sodium cyanoborohydride in 40% ethanol (final concentration). The reaction was incubated at 50°C for 45 min, followed by addition of 20 mM sodium cyanoborohydride and further incubation for 45 min. The reaction was quenched by acidification with 1% TFA. Supernatants were depleted of undecanal and undecanal-modified peptides using HR-X (M) cartridges (Macherey-Nagel). Briefly, cartridges were activated by 2 ml 100% ACN, washed with 2 ml 2% ACN + 0.1% TFA and samples were loaded once on the cartridge and the flow-through containing dimethyl-blocked terminal peptides was collected. To elute remaining dimethyl-blocked terminal peptides from the cartridge, a second elution with 1 ml 40% ACN + 0.1% TFA was performed and the combined flow-through was evaporated in a SpeedVac to a small sample volume which was desalted and purified by C18 StageTips.

#### Optimizing undecanal removal

In this experiment, the removal of undecanal with 40%/50% ethanol and acetonitrile using three different C18 columns was tested. 412µg, 1650µg, and 8250µg undecanal in 0.1% TFA in 40%/50% ethanol and acetonitrile were spun through fully conditioned StageTip, microspin column, or sep-pak C18 columns respectively. The flow-through was collected in 1.5ml Eppendorf tubes and the volume reduced in a SpeedVac. 10µg HeLa peptides and 1M sodium cyanoborohydride were added to a final concentration of 30mM. The volume was adjusted with 40% ethanol to a final volume of 20µL. The samples were incubated at 37°C for 1h and then measured using the quantitative fluorescent peptide assay. The undecanal calibration curve was constructed from 0µg/µL to 41.3µg/µL.

#### Evaluation of peptide recovery dependency on solvent concentrations

Stage-tips with 4 C18 disks were prepared and conditioned with methanol and 0.1% TFA in water. 10µg HeLa peptides were loaded on StageTips and centrifuged at 1200g. The peptides were sequentially eluted with 40% ethanol, 50% acetonitrile, and 80% acetonitrile and collected in 1.5ml Eppendorf tubes. Then, the samples were dried with speed vac and topped up with water to 10µL. The samples were sonicated before performing colorimetric peptide quantification. Elution with 80% and 100% acetonitrile respectively and initial HeLa peptides were used as controls.

### Mass spectrometry

#### Data-dependent acquisition (DDA)

Pre-HUNTER and post-HUNTER HeLa and clinical samples were analyzed on a Q Exactive HF plus Orbitrap mass spectrometer coupled to an Easy-nLC 1200 liquid chromatography (Thermo Scientific) with a 3cm-long homemade precolumn (Polymicro Technologies capillary tubings, 360OD, 100ID), a 35cm-long homemade analytical column (Self-pack PicoFrit columns, 360OD, 75ID, 15um tip ID) and packed with Dr. Maisch beads (ReproSil-Pur 120 C18-AQ, 3um) with a flow rate at 300nL/min and constant temperature at 50°C. Mobile phase A (0.1% formic acid in water) and mobile phase B (0.1% formic acid in 95% acetonitrile) were used for a 65min gradient (3-8%B in 3 min, 8-27%B in 37 min, 27-42%B in 12 min; 42-100%B in 13min). DDA: A full-scan MS spectrum (350-1600 m/z) was collected with resolution of 120,000 at m/z 200 and the maximum acquisition time of 246 ms and an AGC target value of 1e6. MS/MS scan was acquired at a resolution of 60,000 with maximum acquisition time of 118 ms and an AGC target value of 2e5 with an isolation window of 1.4 m/z at Orbitrap cell. The top 12 precursors were selected. Normalized collision energy (NCE) was set to 28. Dynamic exclusion duration was set to 15sec. Charge state exclusion was set to ignore unassigned, 1, and 5 and greater charges. The heated capillary temperature was set to 275°C. It should be noted that 0.8µg peptides in plasma samples, 1µg peptides in 500K and 1M HeLa post-HUNTER samples and all peptides in 10K, 20K, and 100K HeLa post-HUNTER samples were injected for LC-MS/MS analysis.

*Arabidopsis* leaf and rat brain samples were analyzed on a two-column nano-HPLC setup (Ultimate 3000 nano-RSLC system with Acclaim PepMap 100 C18, ID 75 µm, particle size 3 µm columns: a trap column of 2 cm length and the analytical column of 50 cm length; ThermoFisher) with a binary gradient from 5-32.5% B for 80 min (A: H_2_O + 0.1% FA, B: ACN + 0.1% FA) and a total runtime of 2 h per sample coupled to a high resolution Q-TOF mass spectrometer (Impact II, Bruker) as described^7^ (Rinschen et al., 2017). Data was acquired with the Bruker HyStar Software (v3.2, Bruker Daltonics,) in line-mode in a mass range from 200-1500 m/z at an acquisition rate of 4 Hz. The Top17 most intense ions were selected for fragmentation with dynamic exclusion of previously selected precursors for the next 30 sec unless intensity increased three-fold compared to the previous precursor spectrum. Intensity-dependent fragmentation spectra were acquired between 5 Hz for low intensity precursor ions (> 500 cts) and 20 Hz for high intensity (> 25k cts) spectra. Fragment spectra were averaged from t stepped parameters, with 50% of the acquisition time manner with split parameters: 61 µs transfer time, 7 eV collision energy and a collision RF of 1500 Vpp followed by 100 µs transfer time, 9 eV collision energy and a collision RF of 1800 Vpp.

#### Data-independent acquisition (DIA)

The samples were resolubilized in 0.1% formic acid and spiked with iRT peptides before analysis on the Q-Exactive HF system (Thermo) described above. For 1 million *HeLa* samples (1µg of protein was injected), a full-scan MS spectrum (350-1650 m/z) was collected with resolution of 120,000 at m/z 200 and the maximum acquisition time of 60 ms and an AGC target value of 3e6. DIA segment spectra were acquired with a twenty-four-variable window format with a resolution of 30,000 with an AGC target value of 3e6, and using 25% normalized collision energy (NCE) with 10% stepped NCE. The stepped collision energy was 10% at 25% (NCE=25.5 - 27.0 - 30.0). The maximum acquisition time was set to “auto”. DIA method for 20,000 *HeLa* samples was slightly adjusted to accommodate low complexity samples. A 10- variable window format was applied with a resolution of 60,000 and an AGC target of 3e6. The stepped collision energy (NCE) was 28. A default charge state of 3 was applied for MS2 acquisition scans.

#### Data processing

Raw MS DDA data acquired on the Q Exactive HF were processed and searched with MaxQuant^22^ 1.6.2.10 using the built-in Andromeda search engine. The first search peptide tolerance of 20 ppm and main search peptide tolerance of 4.5 ppm were used. The human protein database was downloaded from UniProt (release 2018_09; 20,410 sequences) and common contaminants were embedded from MaxQuant. The “revert” option was enabled for decoy database generation. For analysis of enriched N termini (post-HUNTER) samples, dimethyl (peptide N-term and K) were selected as fixed modifications whereas oxidation (M), acetyl (N-term), Gln->pyro-Glu, and Glu->pyro-Glu were dynamic modifications. ArgC semispecific free N-terminus digestion with maximum two missed cleavage sites. The label free quantification minimum ratio count was 1. “Match between runs” was only enabled for clinical samples. The false discovery rate for PSM, peptide and protein were set as 1%. Label-free quantification was used to quantify the difference in abundance of N termini between samples. To determine dimethyl labeling efficiency and pullout efficiency from pre- and post-HUNTER samples respectively, oxidation (M), acetyl (N-term), dimethyl (K), dimethyl (N-term), Gln->pyro-Glu, and Glu->pyro-Glu were selected as dynamic modifications. ArgC specific digestion mode was used in the first search and Trypsin/P semi-specific digestion mode was selected in the main search. To calculate pullout efficiencies dimethyl (peptide N-term) was defined as variable modification and to calculate labeling efficiencies both dimethyl (peptide N-term and K) were set as variable modifications.

*Arabidopsis* and rat brain DDA data acquired with Impact II Q-TOF instruments were processed and searched with MaxQuant v.1.6.3.3 using embedded standard Bruker Q-TOF settings that included peptide mass tolerances of 0.07 Da in first search and 0.006 Da in the main search. The *Arabidopsis* and rat protein databases were downloaded from UniProt (*Arabidopsis*: release 2018_01, 41350 sequences; rat: release 2017_12, 31571 sequences) with appended common contaminants as embedded in MaxQuant. The “revert” option was enabled for decoy database generation. Database searches were performed as described above, except that enzyme specificity was set as Arg-C semi specific with free N-terminus also in the first search, heavy dimethylation with ^13^CD_2_O formaldehyde was set as label (K) whereas oxidation (M), acetyl (N-term), heavy dimethyl (N-term), Gln->pyro-Glu, and Glu->pyro-Glu were set as dynamic modifications. Data analysis of the *Arabidopsis* seedling experiment considered duplex dimethyl labeling with light ^12^CH_2_O formaldehyde or heavy ^13^CD_2_O formaldehyde (peptide N-term and K).

DIA was analyzed with Spectronaut Pulsar X (version 12.0.20491.0.21112, Biognosys, Schlieren, Switzerland). First, a spectral library was generated by searching the DIA raw files for samples together with 36 DDA files acquired on 12 high-pH fractions for triplicate HeLa samples in Spectronaut Pulsar. The default settings were applied with the following changes: Digest type was semi-specific (free N-terminus) for Arg C, minimum peptide length = 6. Carbamidomethyl (C) and dimethyl (K) were fixed modifications, while variable modifications consisted of oxidation (M), acetyl (N-term), dimethyl (N-term), Gln->pyro-Glu, and Glu->pyro-Glu. The resulting spectral library contained precursor and fragment annotation and normalized retention times. This was used for targeted analysis of DIA data using the default Spectronaut settings. In brief, MS1 and MS2 tolerance strategy were ‘dynamic’ with a correction factor of 1. Similar setting was maintained for the retention time window for the extracted ion chromatogram. For calibration of MS run precision iRT was activated, with local (non-linear) regression. Feature identification was based on the ‘mutated’ decoy method, with ‘dynamic’ strategy and library size fraction of 0.1. Precursor and protein false discovery rate were 1% respectively. The report generated from Spectronaut was filtered for N-terminal peptides with dimethyl and acetyl modifications.

#### Data and statistical analysis

Data evaluation and positional annotation was performed using an in-house Perl script (muda.pl) that combines information provided by MaxQuant, UniProt and TopFINDer^23^ to annotate and classify identified N-terminal peptides. The script (muda.pl) is publicly available (http://muda.sourceforge.io) and will be presented in detail elsewhere. In short, MaxQuant peptide identifications are consolidated by removing non-valid identifications (peptides identified with N-terminal pyro-Glu peptides that do not contain Glu or Gln as N-terminal residue, peptides with dimethylation at N-terminal Pro), contaminant, reverse database peptides, and non-quantifiable acetylated peptides in multi-channel experiments (no K in peptide sequence to determine labeled channel). For peptides mapping to multiple entries in the UniProt protein database, a “preferred” entry was determined by selecting protein entries where the identified peptide matches position 1 or 2, then manually reviewed UniProt protein entries are favored. If multiple entries persisted, the alphabetically first was chosen by default. For *Arabidopsis* seedling experiments, changes in peptide abundance were tested for significance as previously published^24^ using the LIMMA-moderated t-test as implemented in the R limma package. Abundance changes greater than twofold (log2 <-1 or >1) associated with a p-value <0.05 were considered significant.

Proteins identified in human plasma before and after N termini enrichment were annotated with their previously reported plasma protein concentration^17^. N-terminal peptides identified from mitochondria were compared to recently reported N-termini identified in HeLa cells^20^ and listed in the MitoCarta2.0 database^26^. Cleavage site patterns surrounding identified mitochondrial N-termini or altered protease-generated N termini in the *Arabidopsis* seedlings were visualized as iceLogo (https://iomics.ugent.be/icelogoserver/) and WebLogo (https://weblogo.berkeley.edu/logo.cgi).

#### Label free quantification

For label free quantification muda.pl pre-processed data with peptide intensities determined by MaxQuant, is processed further by eliminating termini with intensity values for <20% of the analyzed samples. Data is median normalized, followed by multiplication by the overall data median and log(10) transformation. Pearson correlations, Coefficients of Variation and LIMMA-moderated t-test p-values are calculated using standard implementations in R or python. To retain sample specific termini, missing values were imputed with values randomly selected from a distribution modeled after the tenth to twentieth percentile of the whole data and down-shifted by a random factor of 50-100 placing imputed values into the very low intensity area of the data. Radar plots display the z-score standardized intensity on the y-axis, and fuzzy c-means cluster membership encoded as the line color. Radar plots and t-distributed stochastic neighbor embedding (t-SNE) followed by fuzzy clustering based on imputed data are used for unsupervised characterization of relationships.

## Acknowledgements

We thank Dr. Amina Kariminia and Dr. Kirk Schultz for providing sorted peripheral blood monocytes and Dr. Hans-Jürgen Bidmon for providing rat brains. This work was supported by grants of the Michael Cuccione Foundation (to P.F.L.), the BC Children’s Hospital Foundation (to P.F.L.) and a starting grant of the European Research Council with funding from the European Uniońs Horizon 2020 program (grant 639905, to P.F.H.). P.F.L. is supported by the Canada Research Chairs program and the Michael Smith Foundation for Health Research Scholar program.

## Author contributions

S.W., F.D., S.D., D.T. developed and optimized the protocol. S.W. performed HUNTER analysis on sorted cells. J.T. and S.W. established automation. L.N. and J.T. performed and analyzed automated N termini analysis in patient plasma and bone marrow fluid. A.U. optimized, performed, and analyzed DIA experiments. A.U. and L.N. designed, performed and analyzed mitochondria experiments. F.D. performed protease substrate profiling. F.D., E.K.E. and S.W. performed data analysis. F.D., P.F.H., P.F.L conceived and designed the project. P.F.H. and P.F.L. supervised the project and wrote the manuscript with input from all authors.

## Supplementary Figures

**Fig. S1.**
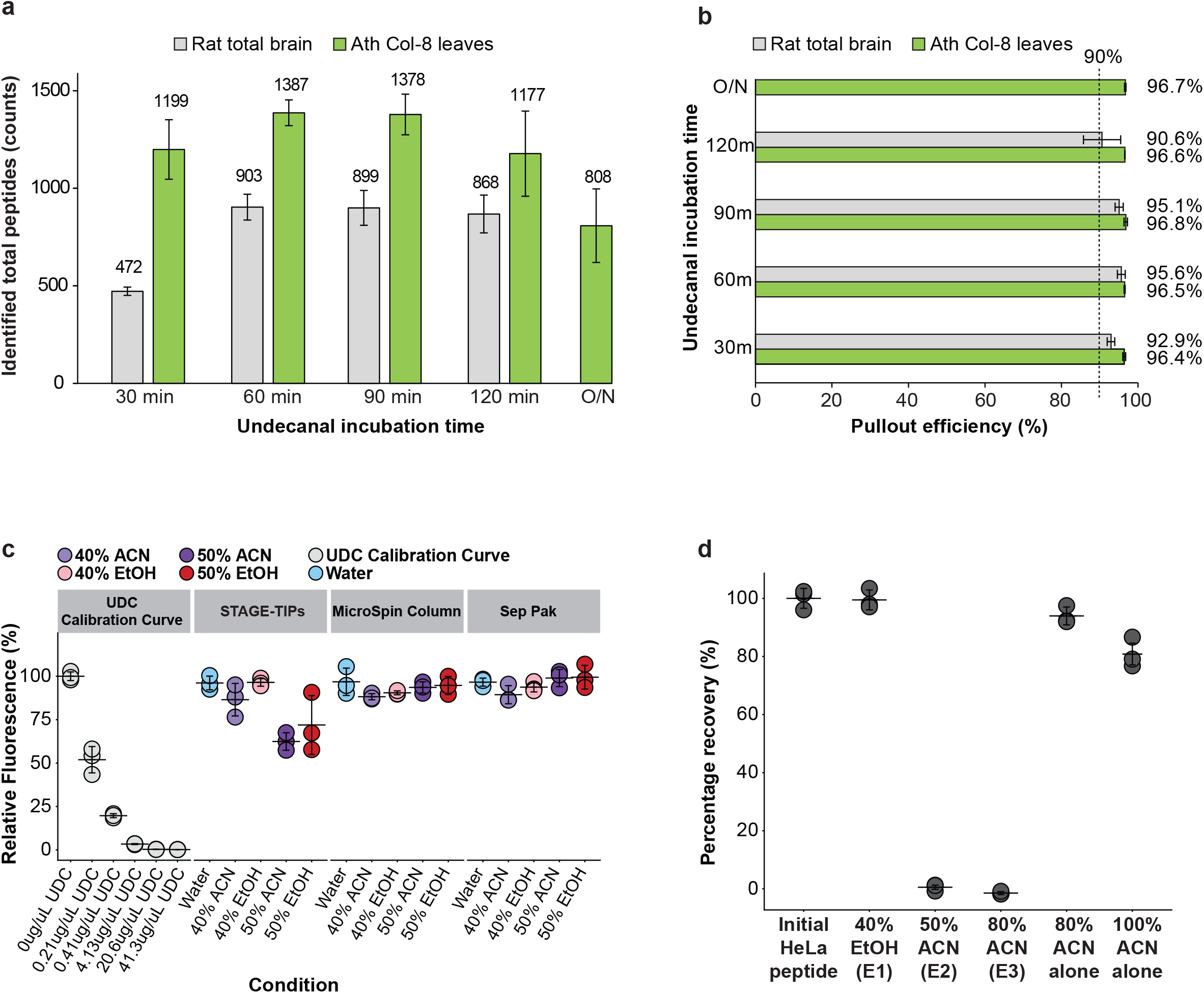
Workflow optimization I – Hydrophobic alkylation reaction time and C18 removal. (**a,b)** Effect of incubation time for undecanal modification of peptides on the number of identified N-terminal peptides. (**a**) and pullout efficiency. (**b**) 200µg of rat total brain and *Arabidopsis* Col 8 whole leaf lysates are used as starting material. Mean of n=3 biological replica, error bars indicate SD. (**c**) Effect of different C18 cartridges and mobile phases (EtOH and ACN) on the depletion of free undecanal. Eluent is incubated with tryptic peptides followed and measured using an amine reactive quantitative fluorometric peptide assay to determine the relative amount of free primary amines on tryptic peptides after reaction with undecanal from the eluent. The UDC calibration curve uses free undecanal. n=3 technical replicates, error bars indicate SD. (**d**) Evaluation of peptide recovery from C18 stage-tips using 40% EtOH as mobile phase. Recovery measured by the amine-reactive colorimetric peptide assay relative to the starting material. In a sequential elution using 40% EtOH followed by 50% ACN and 80%. Tryptic peptides de-salted on C18 using 60% ACN were used as starting material. Mean of n=3 technical replicates, error bars indicate SD.

**Fig. S2.**
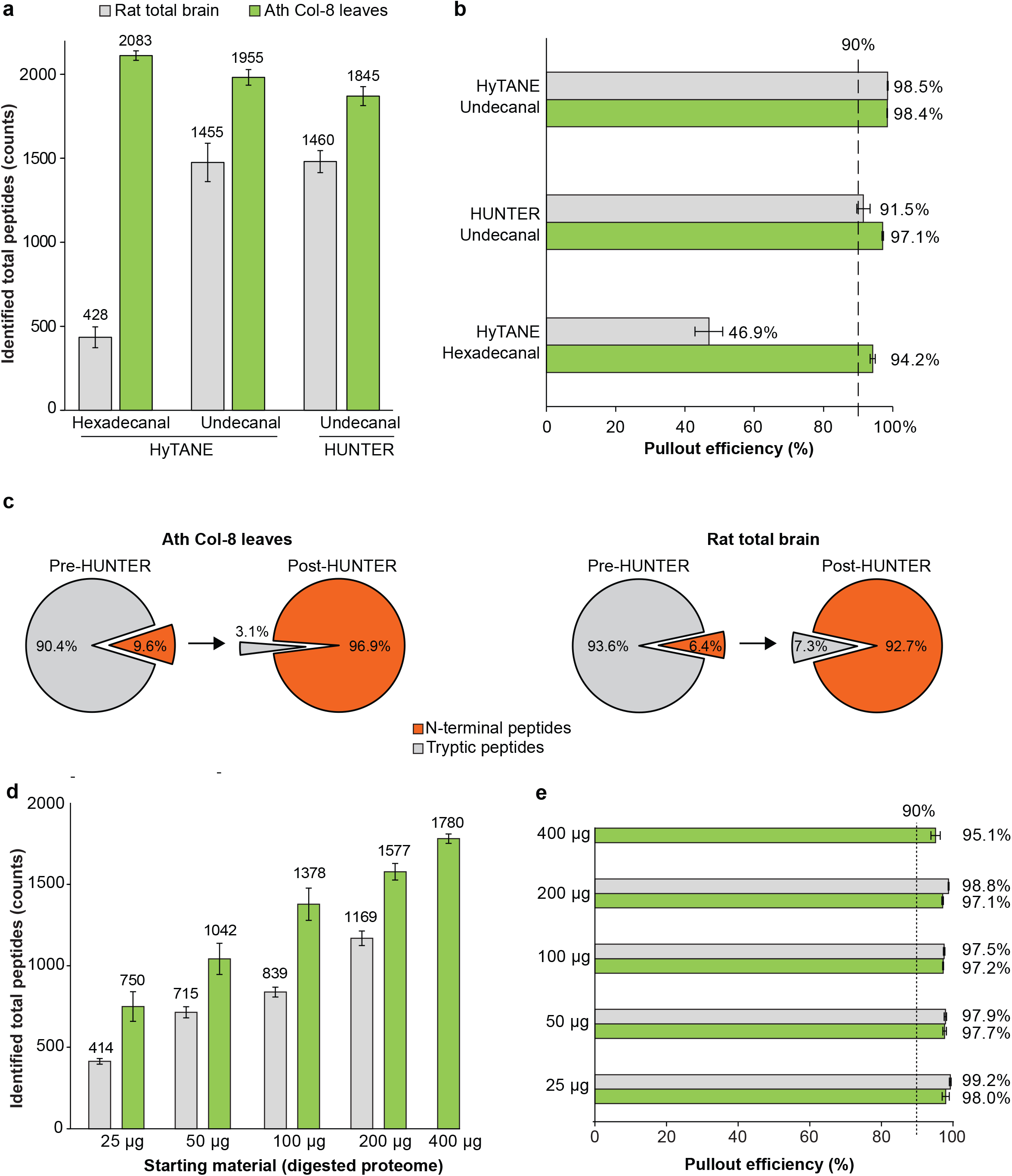
Workflow optimization II – Fatty aldehyde chain length and input amount. (**a**) Peptides identified with adapted hexadecanal or undecanal-based HyTANE protocols with removal of organic solvent after tagging before C18-mediated depletion and after direct depletion of tagged peptides in the presence of organic solvent as implemented in HUNTER. (**b**) Enrichment efficiency assessed as N-terminally modified peptides compared to number of digest-generated peptides with free α-amine. (**c**) Fraction of N terminal and digest-generated peptides in *Arabidopsis* whole leaf and rat total brain proteome digests before (pre-HUNTER) and after (post-HUNTER) enrichment. Shown are average values from three independent biological replicates. (**d, e**) Number of peptides and fraction of N-terminal peptides identified from different amounts of tryptic peptides used for undecanal modification. t=90 min undecanal modification. (**a, b, d, e**) Mean of n=3 biological replicates, error bars indicate SD. A maximum of 1 µg enriched peptides was analysed by nano-LC-MS/MS.

**Fig. S3.**
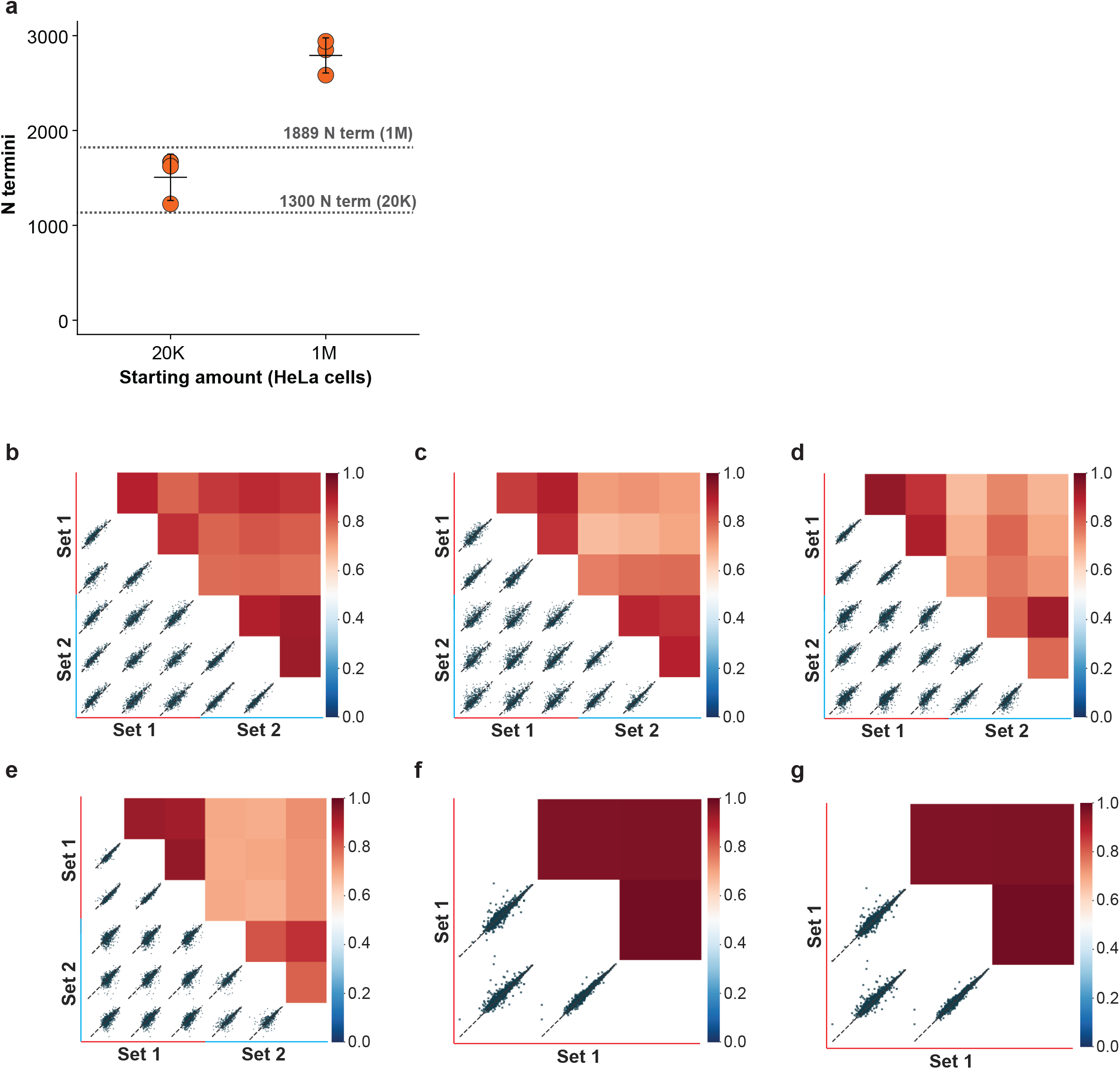
Workflow optimization III – Reproducibility. (**a**) Number of N termini identified from 20K and 1M HeLa cells using DIA. Spectral library built on DDA analysis of 12 high-pH fractions from 1M HUNTER-enriched HeLa cells. Dashed lines show the average number of N termini identified from 20K or 1M HeLa cells by DDA. Mean of n=3 technical replicates. (**b**-**e**) Pearson-correlation between two manually handled HUNTER sets on 10K (**b**), 100K (**c**), 500K (**d**) and 1M (**e**) HeLa cells by DDA. The correlation is based on peptide intensity, n=3 technical replicates. (**f**,**g**) Pearson-correlation between three manually handled HUNTER replicates from 20k (**f**) and 1M (**g**) HeLa quantified by DIA. The correlation is based on sum of elution groups (precursors).

**Fig. S4.**
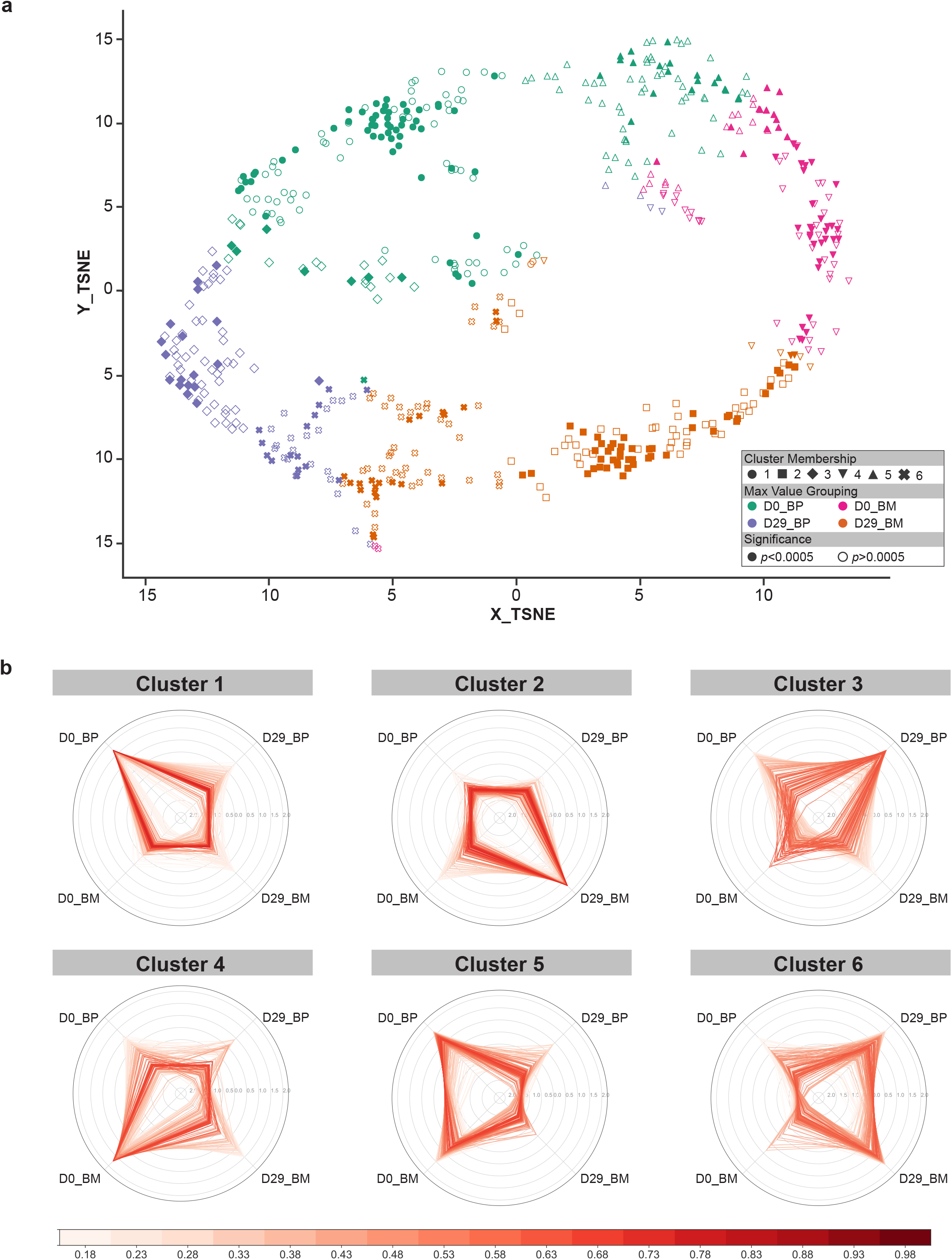
Automated HUNTER analysis of blood plasma (BP) and bone marrow interstitial fluid (BM) form three pediatric B-ALL patients at diagnosis (D0) and after induction chemotherapy (D29). (**a**) Unsupervised t-distributed stochastic neighbor embedding (t-SNE) plot for individual quantified N termini split into 6 clusters (symbols) by fuzzy c-means. The termini are colored based on the condition in which they showed the highest intensity. N termini found to be significantly different (p < 0.0005, ANOVA) are highlighted with solid color. (**b**) Radar plot representation of N termini assigned to the 6 clusters identified in (a). Radial axis represents N termini z-score. Color scaled by fuzzy c-means cluster membership score.

**Fig. S5.**
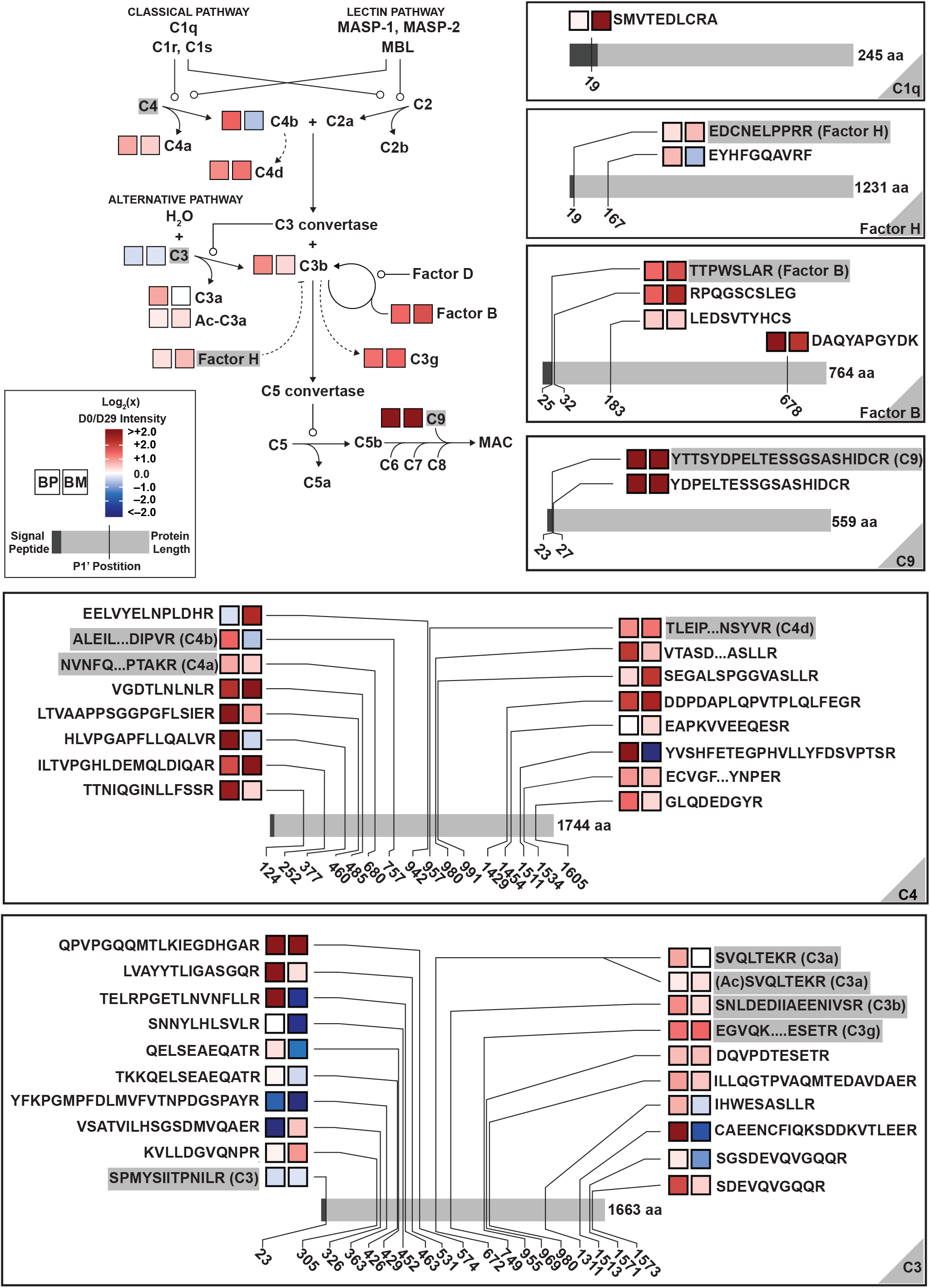
Regulation of the complement pathway in blood plasma (BP) and bone marrow interstitial fluid (BM) of three pediatric B-ALL patients at diagnosis (D0) and after induction chemotherapy (D29). Termini are displayed according to their position on the genome encoded sequence of proteins involved in the complement pathway. Complement proteins are known to be proteolytically cleaved and several fragments with defined start sites are known. Termini matching these start sites are highlighted by grey background and also added to the pathway diagram. Log2 fold changes of termini abundance at diagnosis (D0) and after induction chemotherapy (D29) are color coded from blue (<-2) to red (>2) and visualized for blood plasma (BP) and bone marrow interstitial fluid (BM) next to the respective terminus identified by HUNTER.

**Fig. S6.**
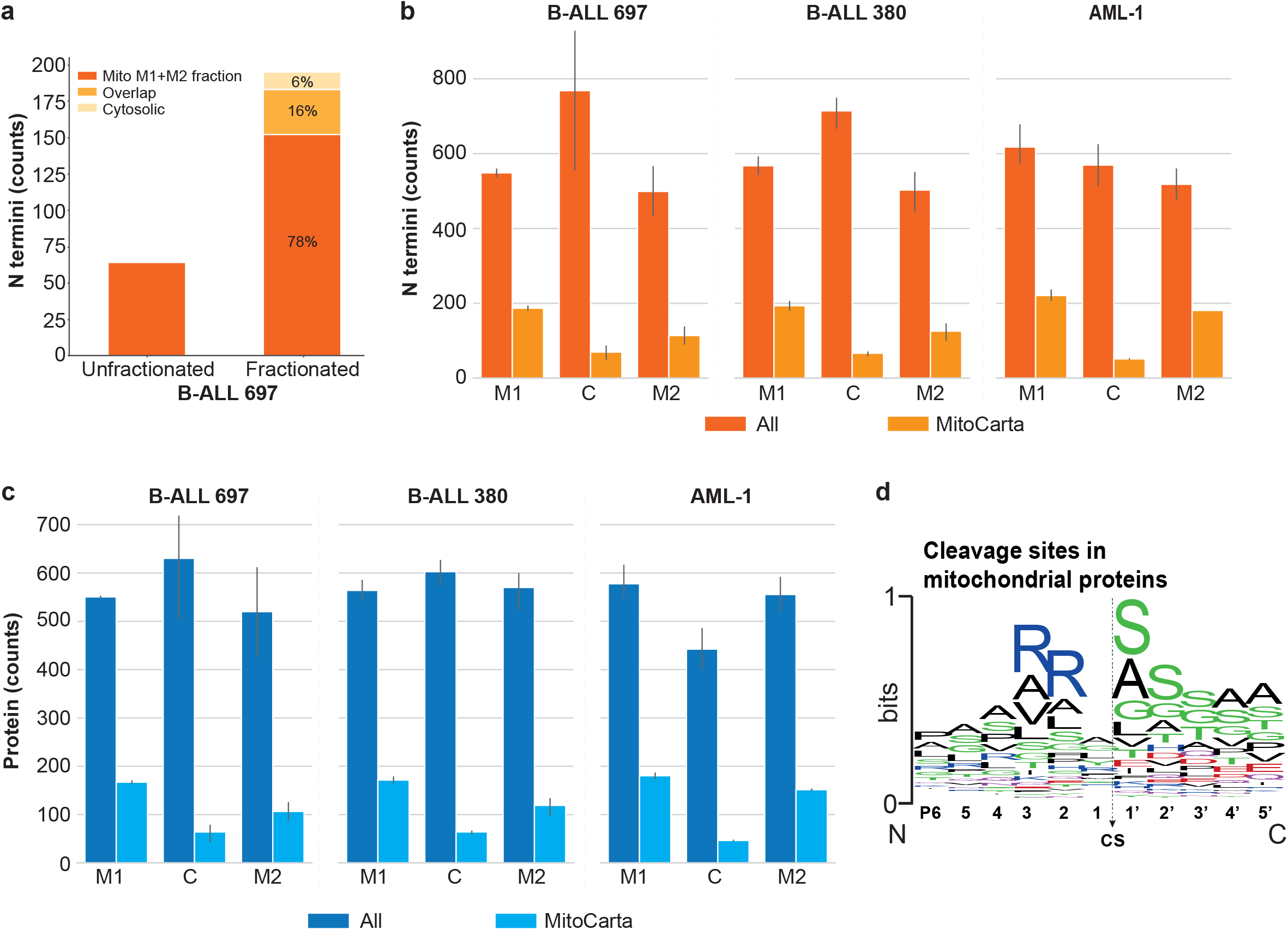
Mitochondrial N terminome from 2.5 million cells by HUNTER. B-ALL cell lines 697 and 380 and BM mononuclear cells from a pediatric AML patient (AML-1) are processed by pressure cycling technology (PCT) supported lysis and fractionated into one cytosolic (C) and two mitochondrial fractions (M1, M2) that are combined for analysis in Fig 2e. (**a**) Total number of identified mitochondrial termini before and after crude subcellular fractionation and enrichment of mitochondria using PCT. After fractionation mitochondrial proteins are identified in the mitochondrial (M1+M2), cytosolic (C) or both fractions. (**b**) N termini identified in each fraction. N termini annotated as mitochondrial by the MitoCarta resource are displayed in light orange. n=2-3, error=SD. (**c**) The total number of proteins identified by N termini in each fraction. Proteins annotated as mitochondrial by the MitoCarta resource are displayed in light blue. n=2-3, error=SD. (**d**) Consensus sequence logo of 233 mitochondrial N termini identified from AML-1 patient cells. The logo resembles the one expected following processing of the mitochondrial import sequence.

**Fig. S7.**
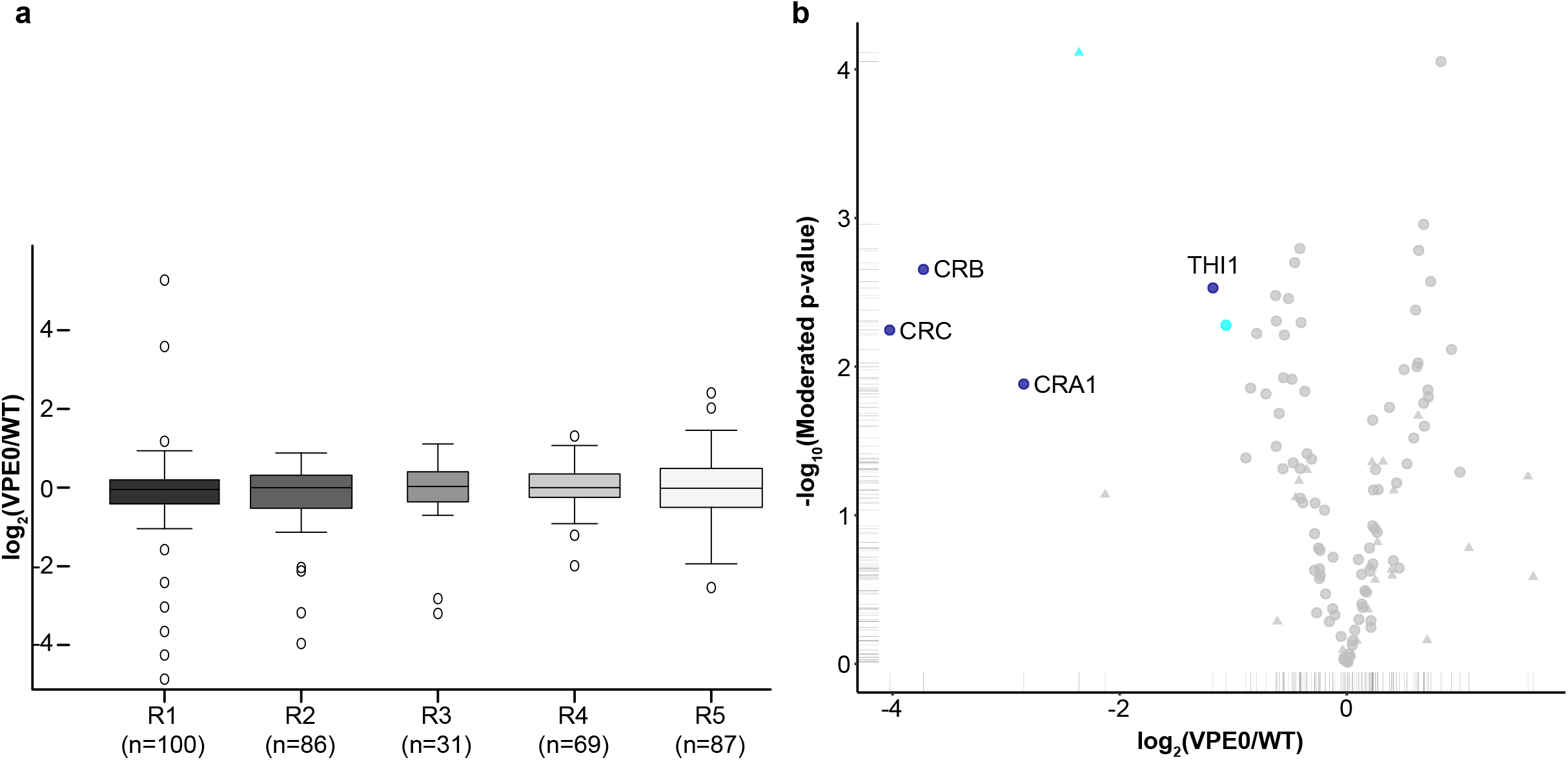
Identification of substrates for seed storage proteases from *Arabidopsis* seedlings. Identification and quantification of protein N termini in single *Arabidopsis* seedlings. (**a**) distribution of log2 ratios in VPE0/wt *Arabidopsis* single seedling HUNTER experiments. (**b**) N termini alterations in 5d old *Arabidopsis*, single VPE0 quadruple mutant/WT seedlings: mainly 12S seed storage proteins (CRA1, CRB, CRC) and THI1 (Thiamine thiazole synthase, chloroplastic) are alternatively processed.

**Fig. S8.**
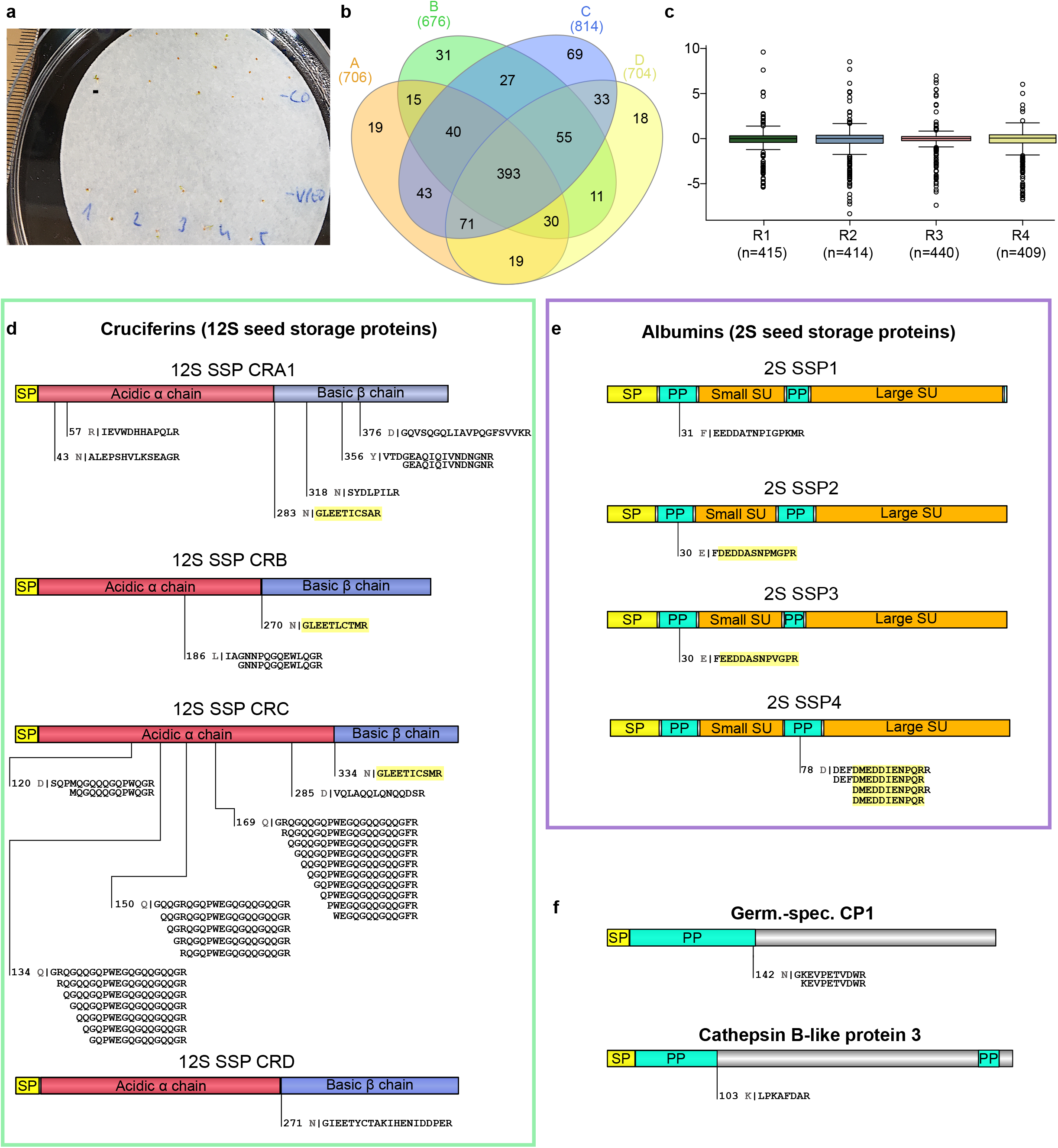
Identification and quantification of protein N termini in *Arabidopsis* seedlings. (**a**) Picture of 2.5d old *Arabidopsis* Col-8 wildtype control (CO) and VPE0 quadruple mutant (VPE0) seedlings, grown on filter paper. Scale bar = 1 mm. (**b**) Venn diagram of protein N termini identified by HUNTER with proteomes extracted from three VPE0 and three Col-8 seedlings per experiment, n=4. (**c**) distribution of log2 ratios in VPE0/wt). (**d**) Significantly depleted processing sites in the VPE0 quadruple mutant seedlings for each of the four major cruciferins (as determined by the moderated t-test acc. to the R package limma): starting position and preceding amino acid (grey) of each N terminal peptide is given. Yellow shaded peptides have already been identified in previous publications (SP: signal peptide). (**e**) significantly enriched proteolytic processing sites in WT seedlings for four 2S seed storage proteins. Yellow shaded peptides have already been identified previously (PP: propeptide, SU: subunit). (**f**) Two examples of significantly enriched pro-peptide proteolytic processing sites in two proteases.

